# First record of anti-predator behavior in the gall-forming aphid *Mordwilkoja vagabunda*

**DOI:** 10.1101/2022.02.16.480690

**Authors:** Andrew Wesley Legan

**Affiliations:** W305 Mudd Hall, Cornell University Dept. of Neurobiology and Behavior, Ithaca NY

**Keywords:** aphid behavior, collective behavior, collective twitching and kicking response CTKR, fortress defense, gall-forming aphid, social insect

## Abstract

The gall-forming aphid *Mordwilkoja vagabunda* has been an outgroup in molecular studies of the evolution of social behavior in the *Pemphigus* genus, but *Mordwilkoja* aphids have not previously been assayed for social behavior, such as altruistic defense. This study reports experiments carried out in July in Ithaca, NY, USA, in which nymphs (immature aphids) of *M. vagabunda* were video recorded under a stereo microscope while they encountered pyralid moth larvae in a plastic arena. *M. vagabunda* nymphs of all instars used their legs to claw moth larvae while pressing their rostrums against the larvae, possibly to pierce the cuticle. Many of the attacking aphids were alatoid nymphs, rather than the specialized first instar soldiers typically observed in *Pemphigus* species. *M. vagabunda* nymphs moved in bursts that sometimes became synchronized among several aphids in the same vicinity. These synchronized, rhythmic movements may be anti-predator defense strategies comparable to the collective twitching and kicking response observed in colonies of *Aphis nerii* and other aphid species. Defensive behaviors by *M. vagabunda* nymphs may be altruistic fortress defense strategies which maximize inclusive fitness of the clone.

**Open Research statement:** Video data are shared publicly on a repository, Zenodo, at this DOI: https://doi.org/10.5281/zenodo.5636845

## Introduction

The independent evolution of sociality in aphids has brought about a diverse array of fascinating behavioral adaptations. Altruistic behaviors of gall-forming aphids include colony defense (“soldier” behavior), gall-cleaning, intergall migration, gall repair, and blocking behavior to obstruct openings in the gall (Stern & Foster, 1996; Pike & Foster, 2008). Shibao et al. (2021) discovered age-dependent task switching between some of these behaviors in *Tuberaphis styraci* (Matsumura). In addition to behavioral adaptations, aphid altruistic behavior may be accompanied by physiological adaptations like venom and hyperclotting excretions (Kutsukake et al., 2004; Kutsukake et al., 2008; Lawson et al., 2017; Kutsukake et al., 2019). Documentation of altruistic behaviors, or lack thereof, in unexamined aphid species sets the stage for future analyses of the evolution of aphid social behavior.

A useful framework for conceptualizing the evolution of social behavior in aphids is outlined by Abbot (2015), who described two key types of evolutionary transitions that together give rise to social aphids. A developmental transition describes the evolution of distinct developmental trajectories for altruistic soldiers, which in social species are sterile and usually frozen during early development, compared to reproductive individuals, which develop to become reproductive adults (Abbot, 2015). A chemosensory transition describes the evolution of the aphid’s ability to detect and respond aggressively to chemosensory cues of predatory insects, which is interesting because this ability has evolved from an herbivorous ancestral state (Abbot, 2015). The timing and degree of these transitions can be better understood by examining behavior of phylogenetically diverse gall-forming aphid species that vary in their social behaviors. *Mordwilkoja vagabunda* (Walsh) is the only described species in the genus *Mordwilkoja*, and because no previous studies have examined behavior in this species, it is unknown whether *M. vagabunda* aphids exhibit altruistic social traits such as anti-predator behavior (Stern & Foster, 1996).

*Mordwilkoja vagabunda* forms irregularly shaped galls (Figure 1) on the stipules of primary host *Populus deltoides* and some other poplar species (Ignoffo & Granovsky, 1961a; Smith, 1971; Blackman & Eastop, 1994; Blackman & Eastop, 2006). The fundatrix produces about 800 offspring which molt through four instars before developing wings as alates (Ignoffo & Granovsky, 1961a). Coincidental with winged alate appearance, the gall develops holes through which alates exit and migrate to secondary host *Lysimachia* spp. (Ignoffo & Granovsky, 1961b; Smith, 1971; Blackman & Eastop, 1994; Blackman & Eastop, 2006). In the fall, sexuals migrate to cottonwoods, where they lay eggs in and on the gall (Ignoffo & Granovsky, 1961a). Eggs overwinter there and give rise to spring fundatrices (gall founders) (Ignoffo & Granovsky, 1961a; Blackman & Eastop, 1994).

**Figure 1:**
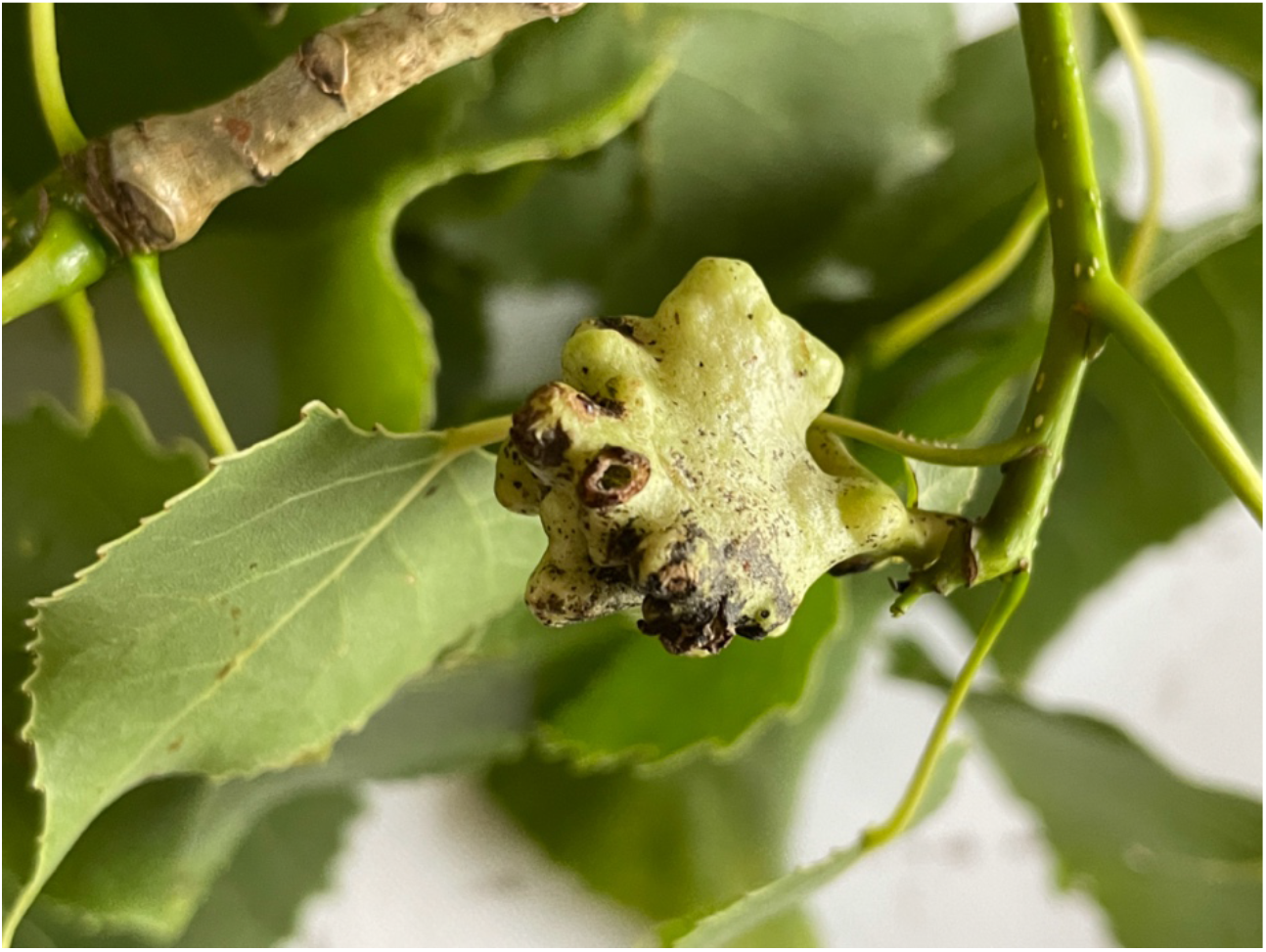
Representative gall of *Mordwilkoja vagabunda* (Walsh).

Pyralid moth larvae are natural predators of some gall-forming aphid species and have been used in experiments to elicit aphid defensive behavior (Aoki & Kurosu, 1992; Stern & Foster, 1996; Shibao & Fukatsu, 2003; Shibao et al., 2010). Tactile stimulation (e.g., gentle prodding using forceps or a fine brush) can also elicit anti-predator defenses from aphids (Sakata & Itô, 1991; Stern & Foster, 1996; Hartbauer, 2010). In this study, *M. vagabunda* nymphs were presented with pyralid moth larvae and experimental probes to elicit anti-predator defenses, which were video recorded.

## Methods

Galls of *Mordwilkoja vagabunda* (Walsh) were identified based on their irregular gall morphology, and were collected from *Populus deltoides* at two sites in Ithaca, NY (site 1: 42°29’43.8”N, 76°27’06.5”W; site 2: 42°29’12.1”N, 76°25’44.2”W). To investigate defensive behavior of *M. vagabunda*, pyralid moth larvae collected from a bumble bee nest box were combined with the contents of *M. vagabunda* galls (3 galls in 3 wells of a 24-well plate). Behavioral assays were conducted in a 24-well plate because the irregular shape of the gall made it difficult to observe and video aphid behavior under the stereo microscope, but it should be noted that removing aphids from the gall and performing experiments under bright light may have altered the aphid’s behavior. It is difficult to distinguish species of pyralid moths by larval morphology, and the larvae used in this study may have been *Plodia interpunctella, Vitula edmandsii*, or *Aphomia sociella*, based on the adult moths that emerged from the bumble bee colony. If the moth larva climbed the wall of the arena, then it was returned to the base of the well using a paint brush. To investigate defensive responses of aphids on the exterior surface of the gall, a paint brush and a metal probe were used to gently prod aphids on the gall surface. Experiments took place between July 6 and July 18, 2021, in Ithaca, NY. Experiments were conducted under a dissecting microscope (Nikon SMZ 745). Microscope magnification ranged continuously between 6.7x and 50x. All experiments were video recorded using an iPhone 8 fixed to an iPhone adapter to microscope 30mm WF lens (iDu Optics). A video summary is provided as supplement (Video S1), and longer videos of experiments are publicly available on Zenodo (DOI: 10.5281/zenodo.5636845). Voucher specimens of aphids and experimentally introduced moth larvae from trials 1 and 2, as well as a collection of 5 galls from site 2 were deposited at the Cornell University Insect Collection (CUIC000051992, CUIC000051993, CUIC000051994; see Appendix S1: Table S1).

## Results

When presented with moth larvae in a plastic arena, *M. vagabunda* nymphs responded by mounting, piercing, and clawing the larvae (Figure 2a,c,d,e,f). After mounting the larvae, aphids moved in bursts as they had been observed doing while on the gall surface. Upon piercing the larva with its stylet, an aphid would move its legs apparently to scratch the larva or else remain still while the stylet remained pressed into the larva. Intermittent movement of mounted aphids sometimes became synchronized, so that the mass of aphids appeared to shimmer in unison (e.g., Video S1a). This pattern of movement appeared to result from the shaking of the aphids’ legs while remaining in the same position on the moth larva. When disturbed by physical contact with experimental probes, *M. vagabunda* nymphs on the gall surface moved with a rhythmic gait while either approaching the probe or raising abdomens and back legs into the air towards the probe. In some cases, the nymphs mounted the paint brush (Figure 2b; Video S1d). Their movement continued in regularly timed bursts until the paintbrush was removed, after which the aphids gradually reduced their movement until they were still. In the event of an encounter between two nymphs on the gall surface, the bursts of movement eventually became synchronized, and then the nymphs walked away from each other (e.g., Video S1d).

**Figure 2:**
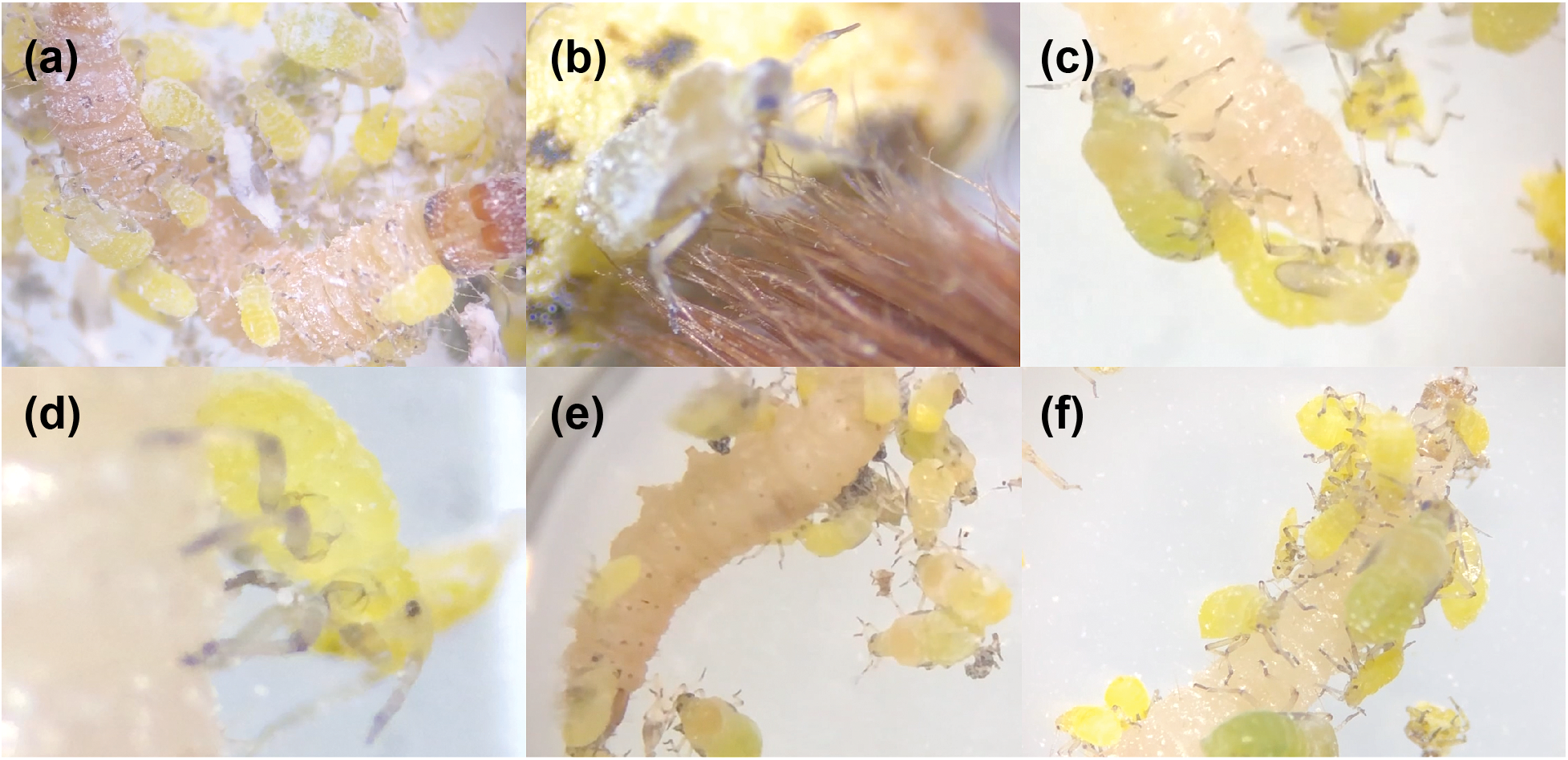
*Mordwilkoja vagabunda* nymphs display risky anti-predator behavior. (a) *M. vagabunda* nymphs of various ages attack a moth larva (from Video S1b). (b) On the external surface of a gall, an *M. vagabunda* nymph (fourth instar) mounts and probes a brush with stylet (from Video S1d). (c) Two fourth instar *M*. *vagabunda* nymphs probe a moth larva with their stylets, and younger aphids attack the larva anteriorly (from Video S1a). (d) A nymph of early instar grips a moth larva while attempting to pierce the moth’s cuticle with its stylet (from Video S1a). (e) In this encounter, defense was less conspicuous, but there were fewer aphids (Video S1c). (f) *M*. *vagabunda* nymphs of various ages attack a moth larva (from Video S1a). Magnification ranges continuously from 6.7x to 50x. Images are freeze-frames from video files available on Zenodo (DOI: 10.5281/zenodo.5636845).

## Discussion

The purpose of this study was to explore defensive behavior of *Mordwilkoja vagabunda* (Walsh) nymphs in July in Ithaca, NY. This is the first record of anti-predator defensive behavior by *M. vagabunda* nymphs. Often the *M. vagabunda* nymphs that mounted and attacked the moth larvae were alatoid (wing-padded) nymphs. Among examined aphid species, defensive behavior in alatoid nymphs is uncommon (Abbot et al., 2018). However, this has been observed in some aphid species (e.g., *Epipemphigus niisimae* (Matsumura), Aoki et al., 1996; *Dermaphis (Dinipponaphis) autumna* (Monzen), Aoki et al., 1999; *Grylloprociphilus imbricator* (Fitch), Aoki et al., 2001). Alatoid nymph defenders in *M. vagabunda* are unlikely to be sterile soldiers. Therefore the chemosensory transition towards altruistic defense may have outpaced the developmental transition towards sterile castes in this species (Abbot, 2015). Future work should assess *M. vagabunda* defensive behavior within the gall to confirm that the observations reported here represent the aphids’ natural behavior, rather than behavior caused by the removal of aphids from the gall or caused by the bright light emitted by the stereo microscope.

*M. vagabunda* aphids raised and waved their back legs and abdomens in response to tactile stimulation and synchronously moved during encounters with conspecifics on the gall surface, and they collectively twitched in synchrony during defense against moth larvae. These observations suggest the role of vibrational cues in *M. vagabunda* anti-predator behavior. Vibration cues are important in predator detection and avoidance in the pea aphid *Acyrthosiphon pisum* (Harris) (Gish, 2021). Sustained substrate-borne vibrations cause several aphid species to leave the plant altogether (Parent et al., 2021). In addition to perceiving vibrations, aphids produce vibrations through body movement. Synchronous movement of aphids within colonies of *Aphis (Toxoptera) aurantii* (Boyer de Fonscolombe) generates vibrations that may function in defense against predators (Broughton & Harris, 2009). Aphid movements can also produce visual deterrents to predators (Sakata & Itô, 1991). Collective leg-waving in the gall-forming aphid tribe Cerataphidini (Hormaphidinae) may function as a bluff to deter large predators (Aoki & Kurosu, 2010). The oleander aphid *Aphis nerii* (Boyer de Fonscolombe) and the cat’s ear aphid *Uroleucon hypochoeridis* (Fabricius) exhibit a “collective twitching and kicking response” (CTKR) that is an effective defense against parasitoid wasps (Hartbauer, 2010). Synchronous movement of this nature was also observed in aggregations of *Aphis fabae* (Scopoli) (Ibbotson & Kennedy, 1951). Combining these published findings with observations from this study, some hypotheses regarding *M. vagabunda* synchronous movement can be proposed.

Non-mutually exclusive hypotheses explaining the function of *M. vagabunda* synchronous movements include: (1) In the context of defense against larger insects, synchronized movement of aphids may function to dislodge the would-be predator and cause it to fall off the surface of the gall, or if the predator is already inside the gall, to fall into a pocket of the irregularly-shaped gall; (2) In the context of defense against larger insects, synchronized movement of aphids may function to elicit movement from the would-be predator, and many aphids could simultaneously assess whether or not the predator is still a threat by perceiving its movement during the period when all aphids are motionless; (3) In the context of encounters with other aphids, synchronized movement may function in recognition, where aphids accept individuals that successfully synchronize their movements and reject individuals that do not synchronize; (4) Synchronized movement of aphids may be a response of aphids to an unobserved stimulus that was introduced by experimental error. More work is required to generate evidence that would support or challenge these and other hypotheses explaining synchronous movement of *M. vagabunda* aphids.

## Supporting information

AppendixS1

## Acknowledgements

The author thanks Serena Peterson for her help in the field. This work was supported by National Science Foundation Graduate Research Fellowship Program grant number DGE-1650441 to Andrew W. Legan and National Geographic Society Young Explorers Grant “Hidden armies: searching for soldiers in unknown social insect species” to Andrew W. Legan.

## Notes

https://doi.org/10.5281/zenodo.5636845

### Competing Interest Statement

The authors have declared no competing interest.

