## AppendixS1 for "First record of anti-predator behavior in the gall-forming aphid *Mordwilkoja vagabunda*"

**Appendix S1**

Table S1: Details of the voucher specimens deposited at the Cornell U. Insect Collection (CUIC)

| CUIC Number | Contents | Lat., Lon. | Date collected | Research number | Trial number | Video S1 |
|---|---|---|---|---|---|---|
| CUIC000051992 | aphids and pyralid moth larva | 42.495503, -76.451802 | July 6, 2021 | AWL00001 | Trial 1 | Video S1A |
| CUIC000051993 | aphids and pyralid moth larva | 42.486691, -76.428939 | July 10, 2021 | AWL00002 | Trial 2 | Video S1B |
| CUIC000051994 | 5 M. vagabunda aphid galls | 42.486691, -76.428939 | July 10, 2021 | AWL00003 | NA | NA |
