## Supplementary material for "First record of anti-predator behavior in the gall-forming aphid *Mordwilkoja vagabunda*": VideoS1_Legend

**Video S1**: Video summary showing highlights of behavioral assays. (a-c) In each of three trials, contents of one aphid gall were combined with a pyralid larva in a well of a 24-well plate to elicit defensive behaviors. (d) *M. vagabunda* nymphs on the surface of the gall were gently prodded with a brush and a metal probe to elicit defensive behaviors.

Videographer: Andrew W. Legan

Cornell University Dept. of Neurobiology and Behavior, Ithaca NY
